# Restraint of Powassan virus replication by TRIM5α facilitates viral avoidance of antiviral immunity

**DOI:** 10.64898/2026.03.15.711935

**Authors:** Rebecca M. Broeckel, Emily A. Fitzmeyer, Eva Chebishev, Jeffrey G. Shannon, Byron Shue, Adam Hage, Stephanie Spada, Efrosini Artikis, Abhilash I. Chiramel, Kristin L. McNally, Charles L. Larson, Maarit von Kutzleben, Shelly J. Robertson, Danielle K. Offerdahl, Hilary A. Lakin, Marshall E. Bloom, Vinod Nair, Nicholas S. Eyre, Michael R. Beard, Michael E. Grigg, Fadila Bouamr, Sonja M. Best

## Abstract

TRIpartite Motif (TRIM) protein 5 alpha (TRIM5α) is a well characterized cellular inhibitor of lentivirus replication that limits transmission of related viruses between primates. We previously reported that TRIM5α derived from humans and rhesus macaques inhibits replication of orthoflaviviruses belonging to the tick-borne encephalitis virus (TBEV) serocomplex, including TBEV, Kyasanur forest disease virus and Langat virus (LGTV), but interestingly not the tick-borne Powassan virus (POWV). To further characterize the primate TRIM5α and orthoflavivirus interface, we screened TRIM5α variants from representative old- and new-world primates for restriction capacity. TRIM5α from old-world African green monkey, De Brazza’s monkey and chimpanzee demonstrated virus-specific restriction of tick-borne orthoflaviviruses. Efforts to determine why TRIM5α fails to inhibit POWV revealed that our lab stock had acquired a non-synonymous mutation in NS3 that, when introduced into a POWV molecular clone, facilitated virus replication in the presence of all inhibitory primate TRIM5α proteins. Infection of human dendritic cells with TRIM5α-resistant POWV resulted in high early replication and strong induction of interferon responses that limited replication compared with the wild-type virus. Thus, primate TRIM5α functions as a potent cellular barrier to infection with tick-borne orthoflaviviruses that restrains replication to a level that may help avoid early innate immune recognition.

## Introduction

Orthoflaviviruses have a remarkable ability to emerge into new geographic regions driven by enzootic transmission cycles that involve ticks or mosquitoes. Infection of humans can result in severe illness, including encephalitis or acute flaccid paralysis, hemorrhagic fevers, hepatitis, and congenital abnormalities (*1*). Currently, approximately a quarter of the world’s population lives in areas endemic to dengue virus (DENV). In 2014, Zika virus (ZIKV) underwent rapid geographical spread after its introduction into the America’s, with the high frequency of human infections revealing a previously unappreciated health threat in the context of congenital infection. Despite the existence of an effective vaccine for yellow fever virus (YFV), approximately 80,000 fatalities occur each year associated with sylvatic transmission cycles and a continual threat of urban transmission (*2*). The prevalence of tick-borne orthoflaviviruses is also increasing, with the last decade witnessing increased geographic foci of tick-borne encephalitis virus (TBEV) across Europe as well as Powassan virus (POWV) in North America (*3*). Development of effective antiviral therapeutics represents an unmet need, necessitating greater understanding of virus replication cycles, host barriers to infection and determinants of human susceptibility to infection and disease.

TRIpartite Motif-containing protein 5 alpha (TRIM5α) is a cellular restriction factor originally identified as a major determinant of the resistance of primate species to infection with human immunodeficienty virus-1 (HIV-1) (*4, 5*). *TRIM5* encodes the three structural domains characteristic of the TRIM protein family including RING (Really Interesting New Gene), B box and coiled-coil domains. The α isoform encodes a C-terminal PRYSPRY domain (also termed B30.2, SPRY, or B30.2/SPRY domain) that functions as the viral recognition domain. In the context of retroviruses, primate TRIM5α variants can form a hexameric cage around incoming mature viral cores, which is thought to accelerate viral uncoating and impede reverse-transcription of the viral genome (*4*). Association of TRIM5α with viral cores also results in activation of AP-1 (Activator Protein-1) and NF-κB (Nuclear Factor kappa-light-chain-enhancer of activated B cells) transcription factors to stimulate innate immune signaling (*6*), as well as exposure of viral nucleic acids for recognition by cyclic GMP-AMP synthase (cGAS) and stimulator of interferon genes (STING) signaling (*7*).

*TRIM5* genes arose during mammalian evolution (*8*). Positive selection has driven innovation of the primate *TRIM5* gene, particularly in the C-terminal PRYSPRY domain (*9*) which is the primary determinant of retrovirus recognition and primate-species specific restriction capacity (*8*). Evolutionary adaptation of TRIM5α has been driven by distinct episodes of endogenous retrovirus infection and subsequent retrotransposition events prior to the existence of primate lentiviruses like HIV-1 (*8*). As a consequence of these evolutionary events, TRIM5α detection of HIV capsids may be a selective bottleneck, dictating which contemporary zoonotic HIV strains have become widely transmitted in humans (*10*). However, the potential for TRIM5α as a strong barrier and selective pressure for viral evolution outside of retroviruses is unknown.

Until recently, a significant role for primate TRIM5α in restriction of viruses other than retroviruses had not been reported. However, our group showed that endogenous cellular expression of human TRIM5α strongly inhibits replication of orthoflaviviruses belonging to the TBEV serocomplex, including Kyasanur forest disease virus (KFDV; a hemorrhagic fever causing virus endemic to India) and Langat virus (LGTV; a virus with low human pathogenic potential isolated from ticks in Malaysia and Thailand), but interestingly not POWV or any of the mosquito-borne viruses we tested including DENV, ZIKV, YFV or West Nile virus (*11, 12*). RNA replication of these viruses is orchestrated by nonstructural protein 5 (NS5, encoding the methyltransferase and RNA-dependent RNA polymerase) and NS3 (encoding serine protease and RNA helicase functions) in specialized replication organelles derived from the endoplasmic reticulum (ER). TRIM5α binds to, and promotes degradation of, NS3 and its co-factor NS2B. NS2B is required for tethering to the ER and maturation of the NS3 protease active site. Indeed, the TRIM5α PRYSPRY domain recognized NS3 only in the context of a NS2B/3 fusion (*12*). This suggests that, like sensing of the retroviral core, recognition of NS2B/3 by TRIM5α may be dependent on substrate conformation. These initial findings raise several important questions including what determines genetic sensitivity of specific orthoflaviviruses to restriction by TRIM5α, the degree to which TRIM5α variants from diverse primate species can differentially recognize infection by different orthoflaviviruses, and whether viral evasion of TRIM5α has implications for virus replication and the human immune response.

Here, we explore some of the principle features of TRIM5α-mediated restriction previously defined in the context of lentivirus restriction and apply them to infection with orthoflaviviruses. The specific nature of how TRIM5α engages orthoflavivirus NS3 has direct implications for understanding innate immune barriers for vector-borne orthoflaviviruses in primates.

## Results

### Tick-borne orthoflaviviruses are sensitive to restriction by TRIM5α from multiple old-world primate species

TRIM5α from humans and rhesus macaques can inhibit replication of TBEV, KFDV, and LGTV, whereas the TRIM-Cyp fusion protein of pig-tailed macaques, where the PRYSPRY domain has been replaced by a retrotransposition of cyclophilin A, fails to restrict replication of these viruses (*11, 12*). The requirement of the TRIM5α PRYSPRY domain for interactions with orthoflavivirus NS2B/3 (*12*) suggests that positive selection in primate *TRIM5* genes may have resulted in primate-specific restriction capacity towards the orthoflaviviruses. To test this hypothesis, HEK293 cells were generated to stably express TRIM5α derived from old-world (patas monkey, grivet monkey, sabaeus monkey, De Brazza’s monkey and pig-tailed macaque as a control) and new-world (squirrel monkey, marmoset, red howler monkey, and owl monkey) primates as well as a hominid (chimpanzee) (Figure 1A). Following infection, expression of grivet (*Chlorocebus aethiops*) or sabaeus (*Chlorocebus sabaeus*) TRIM5α restricted all three tick-borne viruses tested at 24 hours post infection (hpi) by 10-fold or greater (TBEV, KFDV and LGTV) with the grivet variant derived from Cos-7 cells exibiting stronger inhibition than that from sabaeus Vero cells. The TRIM5α variant derived from De Brazza’s monkey (*Cercopithecus neglectus*) inhibited replication of LGTV and TBEV but not KFDV whereas expression of TRIM5α from chimpanzee (*Pan troglodytes*) lead to a moderate reduction of TBEV replication at 24 hpi but not other viruses tested. No new-world primate TRIM5α sequence tested was inhibitory towards any of the tick-borne orthoflaviviruses. ZIKV was also tested as a representative of the mosquito-borne orthoflaviviruses that uses humans to maintain urban transmission cycles, but its replication was not affected by stable expression of any primate TRIM5α (Figure 1B, C). Together, this work demonstrates that sequence variation in primate *TRIM5* genes greatly influences restriction capacity against tick-borne orthoflaviviruses and seems to limit restriction to old-world primates. However, within old-world primates, TRIM5α-mediated restriction is not limited to one lineage, with primates from both Cercopithecini (De Brazza’s, grivet, sabaeus) and Papionini (rhesus) exhibiting strong restriction phenotypes to the same virus (LGTV and TBEV).

**Figure 1:**
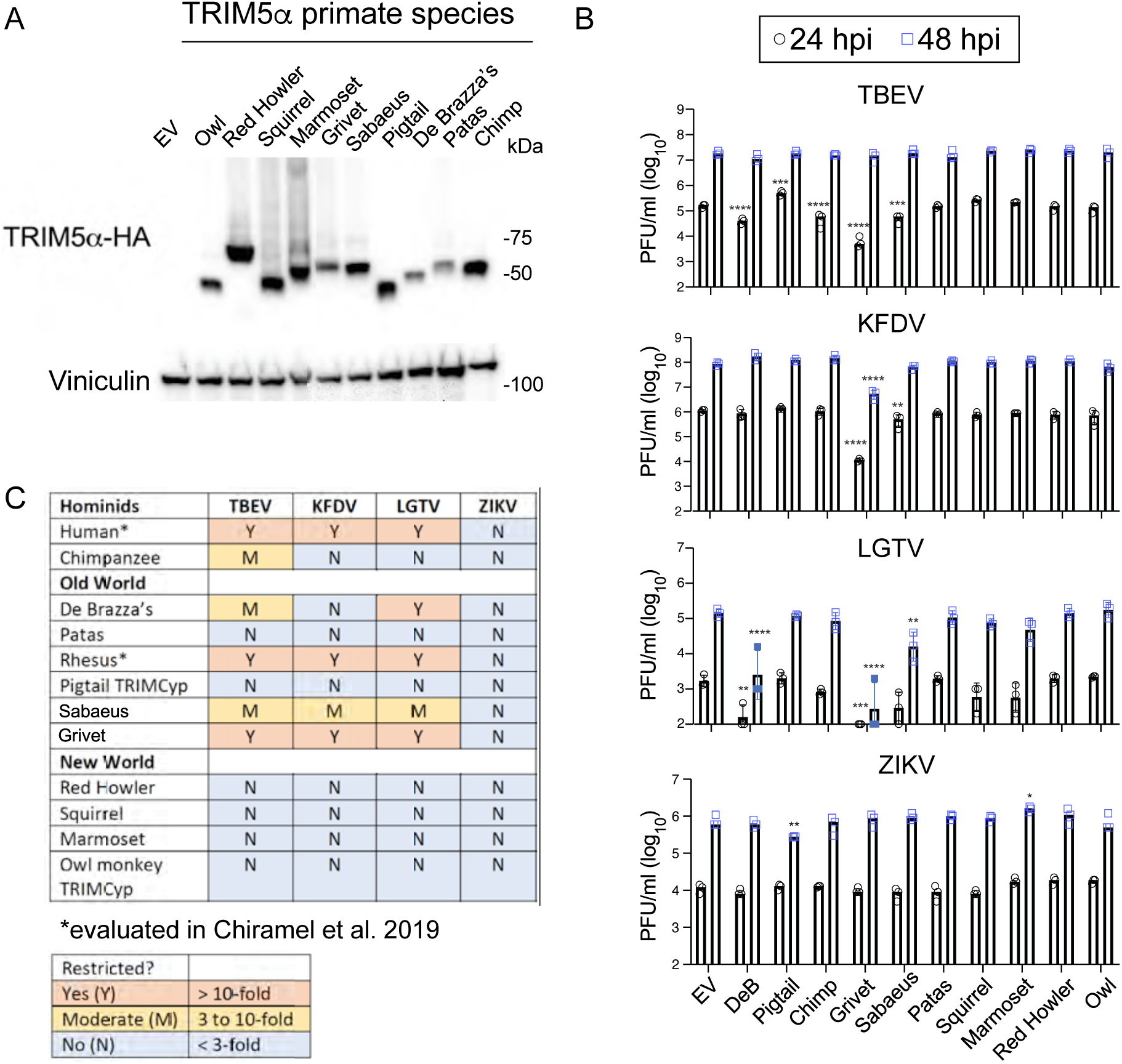
Tick-borne orthoflaviviruses are sensitive to restriction by TRIM5α from multiple old-world primate species. **A.** Western blot demonstrating stable expression of TRIM5α-HA from select old- and new-world primate species in HEK293 cells. **B.** Infectious titers following infection of cell lines shown in **A** at 24 and 48 hpi with tick-borne encephalitis virus (TBEV), Kyasanur Forest Disease virus (KFDV), Langat virus (LGTV) and Zika virus (ZIKV). Values represent the mean ± SD (n=3). Statistical significance was assessed using two-way ANOVA followed by Dunnett’s post hoc test for multiple groups. * p<0.05, ** p<0.01, *** p<0.001, **** p<0.0001. **C.** Summary of restriction sensitivity data in **B**.

### LGTV sensitivity to TRIM5α maps to the helicase domain of NS3

Having established that different primate TRIM5α sequences differentially inhibit orthoflaviviruses belonging to the TBEV serogroup, we next aimed to identify viral sequences that confer sensitivity to TRIM5α-mediated restriction. LGTV has limited virulence potential in humans as evidenced by the trial of LGTV as a live-attenuated vaccine candidate for protection against TBEV infection (*13*). Therefore, we chose LGTV to serially passage in HEK293 cells stably expressing rhesus (rh) TRIM5α. By passage 9, LGTV from duplicate passages replicated to equivalent titers in HEK293-rhTRIM5α cells and HEK293-empty vector control cells, while virus stock passaged in HEK293-empty vector control cells remained sensitive to TRIM5α-mediated restriction, producing ∼1000 fold less titers at 24 hpi (Figure 2A). Virus genome sequencing revealed that compared to the starting stock virus, mutations present in >90% of reads and unique to the two variants passaged in HEK293-rhTRIM5α cells occurred at nucleotide positions T4800C (NS3 V68A) and G6272A (NS3 V559M) (Variant 1; Figure 2B) or T6273C (NS3 V559A) (Variant 2) (Figure 2C). In addition, nucleotides A9735G (NS5 D691G) and T9765A (NS5 F701Y) were present in mixed populations in the stock virus but strongly selected for in both variants passaged in HEK293-rhTRIM5α cells (Figure 2D).

**Figure 2:**
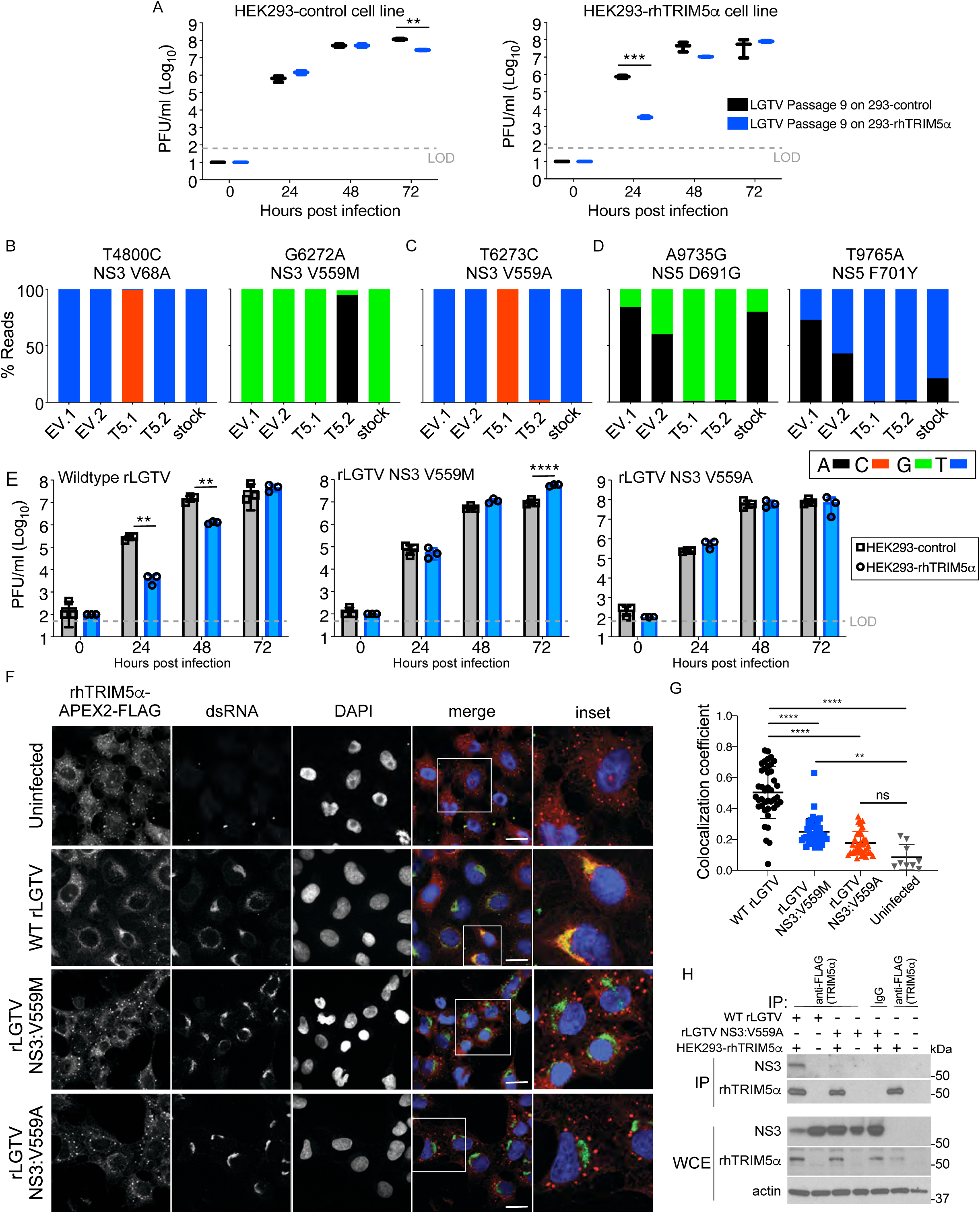
Langat virus sensitivity to inhibition by rhTRIM5α maps to the NS3 residue V559. **A.** LGTV was serially passaged in HEK293 cells stably expressing rhTRIM5α-HA or in a control cell line. At passage 9, growth curves of virus stocks were performed in the same cell lines demonstrating loss of restriction sensitivity of HEK293-rhTRIM5α-passaged virus. Data from 1 of 2 variants from duplicate samples passaged in control or HEK293-rhTRIM5α cell lines is shown. **B-D.** Summary of RNAseq data of LGTV starting stock virus and duplicate passages in HEK-control cells expressing an empty vector (EV) or rhTRIM5α-HA showing non-synonomous substitutions present in >90% of reads and **B.** unique to NS3 of the T5.1 variant, **C.** unique to NS3 of the T5.2 variant, or **D.** commonly enriched in the NS5 of T5.1 and T5.2. **E.** Growth curves of recombinatnt WT LGTV (rLGTV), rLGTV NS3 V559M, or rLGTV NS3 V559A in HEK293 cells stably expressing rhTRIM5α-HA or in control (empty vector) cells (n=3). **F.** Immunoflourescence demonstrating localization of rhTRIM5α-FLAG (red) in HEK293 cells infected with WT or TRIM5α-resistant LGTV variants. Infection status of the cell is confirmed by positive staining for dsRNA (green); DAPI stain of cell nucleus (blue). Scale bars represent 12µm. **G.** Colocalization coefficient of dsRNA and rhTRIM5α-FLAG on a per cell basis (n=9-40 cells). **H.** Co-immunopreciptiation of rhTRIM5α-FLAG and NS3 in cells infected with WT rLGTV or rLGTV NS3 V559A. Data is shown from one of three experiments performed. All graphed values represent the mean ± SD. Statistical significance was assessed using two-way ANOVA followed by Sidak’s post hoc test for multiple groups. * p<0.05, ** p<0.01, *** p<0.001, **** p<0.0001.

The independent selection of mutations G6272A (NS3 V559M) and T6273C (NS3 V559A) that were not observed in the stock virus population strongly suggest that NS3 V559 is important for restriction-sensitivity of LGTV to rhTRIM5α. To test this, we engineered a molecular clone of LGTV (strain TP21) using the circular polymerase extension reaction (CPER) method, modelled on a clone for ZIKV (*14*). Wildtype (WT) recombinant LGTV (rLGTV) exhibited sensitivity to rhTRIM5α-mediated restriction whereas rLGTV substituted at NS3 V559M or V559A was insensitive to restriction. Compared to WT rLGTV, rLGTV-NS3 V559A exhibited no replication deficit, whereas rLGTV-NS3 V559M replicated ∼0.5 log_10_ lower at 24 and 48 hpi (Figure 2E). Together, these data demonstrate that a single amino acid substitution in NS3 at V559 is sufficient to avoid restriction by rhTRIM5α.

Although resistance to rhTRIM5α can be achieved by mutation of NS3 at V559, evasion also appears to select for mutations within the NS5 RNA-dependent RNA polymerase domain suggesting that TRIM5α recognizes NS3 in the replication organelle in the ER. Co-localization between viral dsRNA replication intermediates and rhTRIM5α has been previously demonstrated (*12*). Consistent with previous findings, TRIM5α was recruited from distinct cytosolic puncta to sites co-staining for dsRNA in WT rLGTV-infected cells (Figure 2F). However, TRIM5α remained in cytosolic bodies following infection with rLGTV-NS3 V559A or rLGTV-NS3 V559M, and had significantly less co-localization with dsRNA (Figure 2G). Interaction of TRIM5α with NS3 was reduced following co-immunoprecipitation (co-IP) in rLGTV NS3 V559A-infected cells, despite higher NS3 expression than WT rLGTV due to lack of restriction (Figure 2H). We also visualized TRIM5α localization by transmission electron microscopy (TEM) using HEK293 cells stably expressing a rhTRIM5α-APEX2-FLAG fusion protein. Here, APEX2 catalyzes the polymerization and local deposition of diaminobenzidine that then recruits electron-dense osmium to provide EM contrast to visualize protein localization (*15*). TRIM5α appeared as cytosolic puncta with diffuse definition in uninfected cells (Figure 3). Clear relocalization of TRIM5α to viral replication organelles was observed in cells infected with WT rLGTV but not with rLGTV NS3 V559A. Thus, substitution of NS3 V559 for Ala or Met is sufficient to avoid recognition of the LGTV replication organelle by rhTRIM5α.

**Figure 3:**
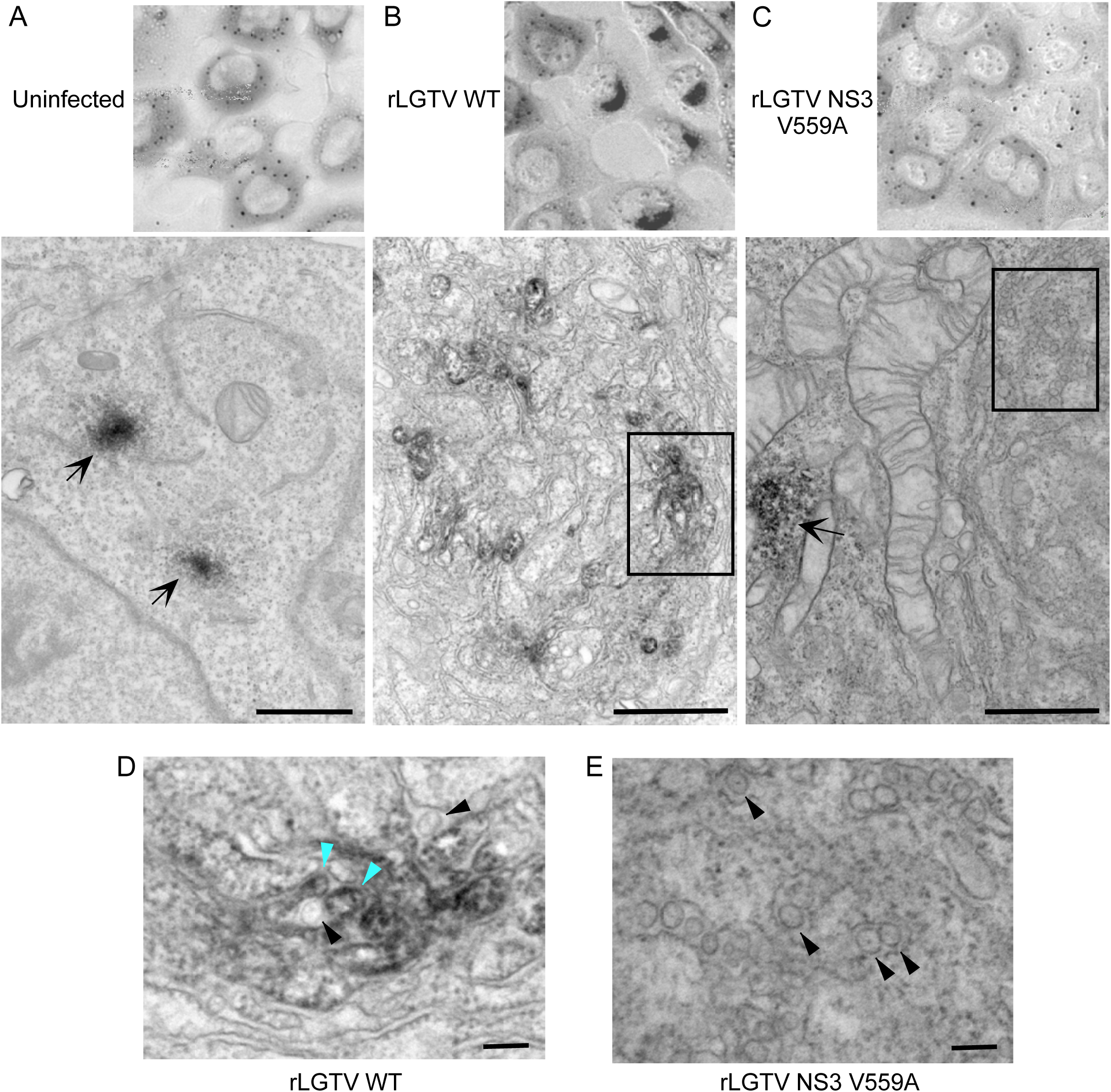
Transmission electron microscopy of rhTRIM5α-APEX2 localization following LGTV infection. HEK293 cells stably expressing rhTRIM5α-APEX2-FLAG were infected with **A.** mock, **B.** WT rLGTV or **C.** rLGTV NS3 V559A and imaged at 48 hpi. Scale bars represent 2µm. Arrows represent cytosolic TRIM5α bodies. **D. and E.** depict increased magnification of areas boxed in B. and C. respecitively. Black arrow heads depict LGTV replication organelles not associated with TRIM5α; blue arrow head depicts TRIM5α association with LGTV replication organelle. Scale bars represent 100nm.

### Resistance of POWV to TRIM5α is a cell culture adaptation to growth in Vero cells

NS3 shares approximately 80% amino acid identity between tick-borne orthoflaviviruses, and Val559 is strictly conserved (Figure 5A). Figure 4 depicts a backbone-atom alignment of the available PDB NS3 helicase structures for LGTV, KFDV and TBEV with the three subdomains indicated (I, II, III). The backbone root mean square deviation of the structures is less than 1Å and most of the structural variation is attributed to loops between the helices. At the top of Domain III, the V559 loop creates a small hydrophobic pocket with hydrophobic interactions between V556, A557 and V559 (LGTV numbering). Additionally, the backbone of V559 forms a hydrogen bond with the sidechain of R562, maintaining the loop structure. When substituting the Val at position 559 for Ala (LGTV Variant 2), the backbone hydrogen bond between V559A and R562 persists and the orientations of neighboring residues T560 and D561 remain similar. The small Ala sidechain is easily accommodated and does not pose a steric hinderance with H534 on the adjacent helix. However, the substitution of V559 to Met (LGTV Variant 1) limits the orientation of the Met sidechain as it lays parallel to the loop avoiding a steric clash with the imidazole ring of H534. In this orientation, the Met may form a Met-aromatic interaction with H534 allowing for loop stabilization. Both of the V559 variants maintain the hydrophobic nature of the region by forming a hydrogen bond with R562, a highly conserved residue throughout orthoflaviviruses (Figure 5A).

**Figure 4:**
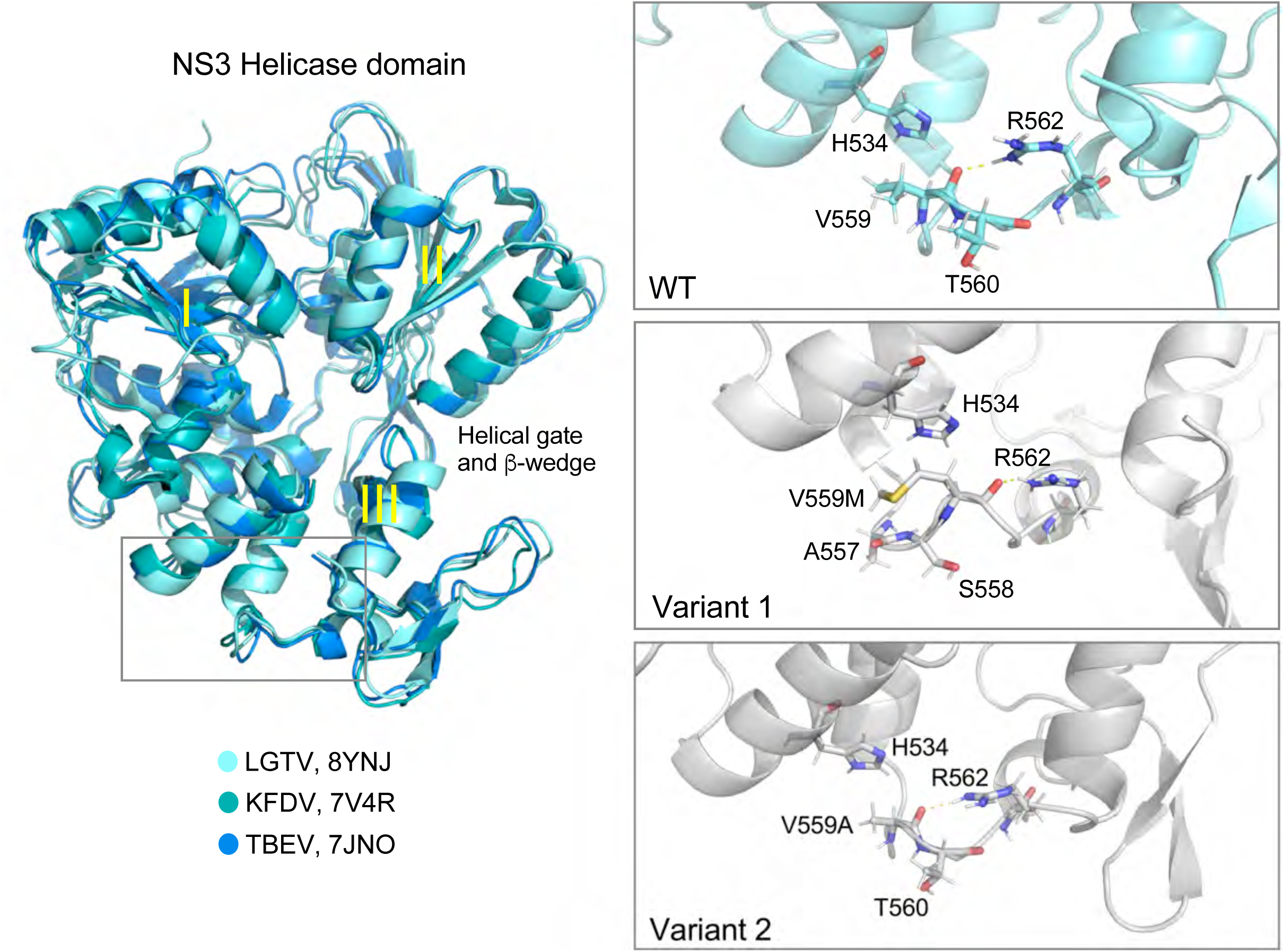
Structural conservation of NS3 helicase domain between tick-borne orthoflaviviruses. PDB structures from LGTV, KFDV and TBEV NS3 helicase are shown. Subdomains I, II, and III and the helical gate and β-wedge are indicated. The V559 loop is boxed in grey and magnified in the right hand panels, with non-synonomous substitions at V559M (Variant 1) or V559A (Variant 2) modeled.

**Figure 5:**
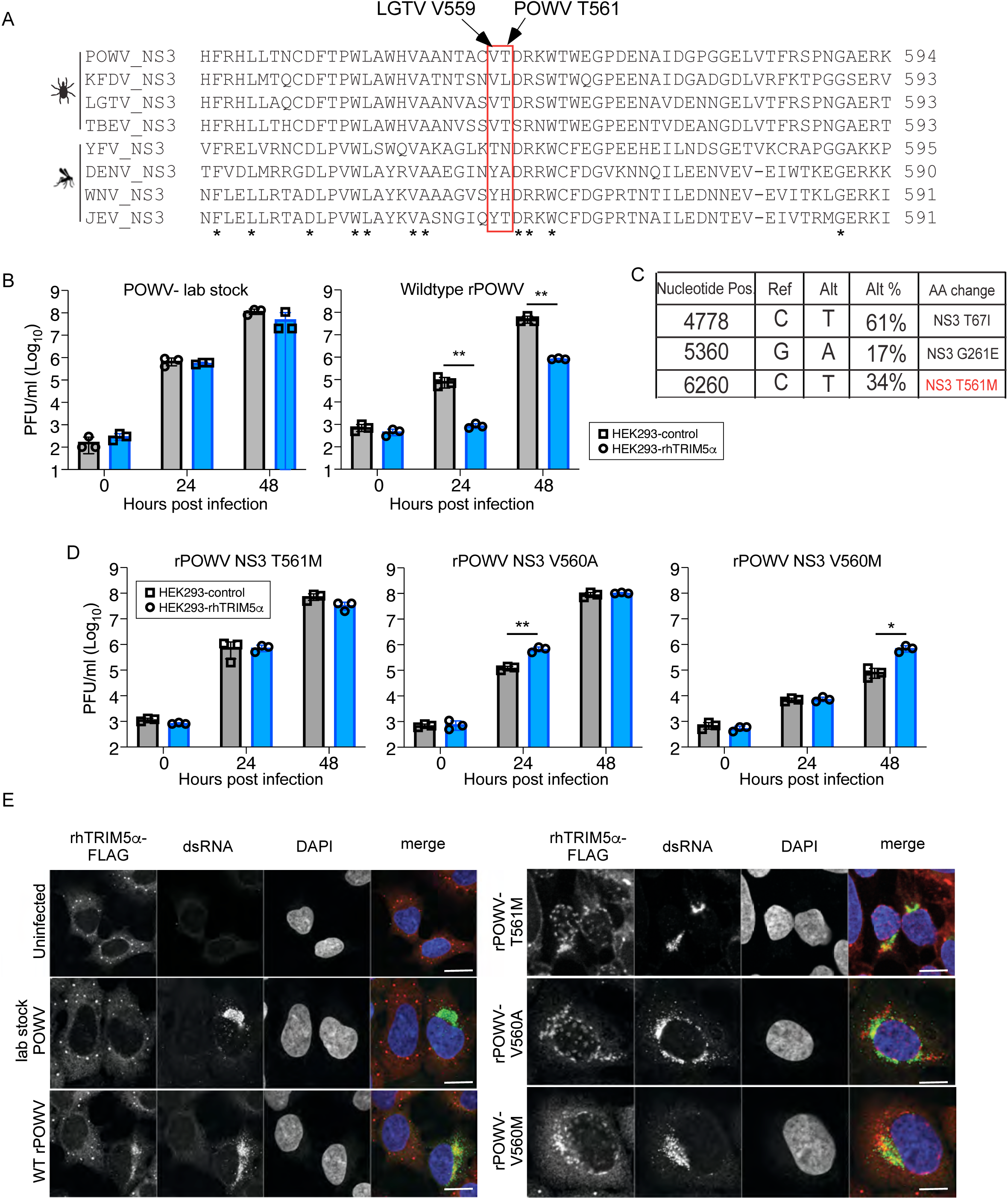
Powassan virus sensitivity to inhibition by rhTRIM5α maps to the NS3 residue T561. **A.** Sequence alignment of orthoflavivirus NS3 residues proximal to LGTV V559. **B.** Growth curves of POWV lab stock or WT recombinant POWV (rPOWV) in HEK293 cells stably expressing rhTRIM5α-HA or in control cells (n=3). **C.** Summary of RNAseq data from the POWV lab stock. **D.** Growth curves of recombinant POWV (rPOWV) bearing mutations at NS3 T561M, V560A or V560M in HEK293 cells stably expressing rhTRIM5α-HA or in control cells (n=3). **E.** Immunoflourescense demonstrating localization of rhTRIM5α-FLAG (red) in HEK293 cells infected with WT or TRIM5α-resistant POWV variants. Infection status of the cell is confirmed by positive staining for dsRNA (green); DAPI stain of cell nucleus (blue). Scale bars represent 10µm. Graphed values represent the mean ± SD. Statistical significance was assessed using two-way ANOVA followed by Dunnett’s post hoc test for multiple groups. * p<0.05, ** p<0.01, *** p<0.001, **** p<0.0001.

The high degree of structural (Figure 4) and sequence conservation (Figure 5A) raises questions as to why POWV was unaffected by TRIM5α in our initial report (*12*). To begin to address this, we generated a POWV molecular clone by CPER (LB strain; GenBank: L06436.1). Replication of our lab virus stock was not affected by ectopic expression of rhTRIM5α, as previously reported. However, titers of recombinant wildtype POWV (WT rPOWV) were inhbited by nearly 100 fold at 24 and 48 hpi (Figure 5B). RNAseq of the lab stock revealed non-synonymous substitutions compared to the reference sequence L06436.1 in NS3 encoding T67I, G261E and T561M (Figure 5C). We therefore tested the T561M mutation due to its location directly adjacent to NS3 V559 (LGTV numbering). Introduction of NS3 T561M alone phenocopied the replication of our lab stock, producing equivalent titers in control and rhTRIMα-expressing cells (Figure 5D). Substitution of the equivalent Val mutation from LGTV at V560A also increased resistance of rPOWV to TRIM5α-mediated restriction, whereas a V560M substitution reduced replication by more than 100 fold even in control cells (Figure 5D), consistent with implications from the structural models. By IFA, TRIM5α did not co-localize with POWV dsRNA in cells infected with our lab stock, whereas colocalization was readily evident in WT rPOWV (Figure 5E). Interestingly, rhTRIM5α partially recognized infection with rPOWV encoding NS3 mutations T561M, V560A, or V560M as it was recruited to perinuclear sites in infected cells, but only partially co-localized with viral dsRNA. Thus, mutation of NS3 T561M confers a high degree of restistance to rhTRIM5α, but in contrast to LGTV, additional structural changes must be required for complete evasion of TRIM5α by POWV.

We hypothesize that TRIM5α resistance of our POWV lab stock was acquired through repeated propagation of stocks in African green monkey Vero cells (*Chlorocebus sabaeus*) that express readily detectable TRIM5α endogenously (Figure 6A). In support of this, rPOWV NS3:T561M replicated to higher titers in parental Vero cells, but replication was equivalent to WT rPOWV in TRIM5^-/-^ Vero cells generated by CRISPR/Cas9 gene editing (Figure 6A). Furthermore, the NS3 T561M and V560A mutations conferred resistance to TRIM5α from African green and De Brazza’s species (Figure 6B) in addition to rhesus macaque (Figure 5D). Mutations in viral proteins present in the replication complex (ie NS3) could drive increased RNA replication to outcompete virus restriction factors through saturation. However, while rPOWV NS3 T561M produced 100-fold more infectious virus than WT rPOWV in parental Hap1 cells, WT and mutant viruses replicated to equivalent titers in TRIM5^-/-^ Hap1 cells (Figure 6C). Thus, in two cell lines, Vero and Hap1, NS3 T561M does not confer a replication advantage unless a restrictive TRIM5α is expressed. These findings suggest that loss of restriction by rPOWV NS3 T561M is due to loss of TRIM5α recognition of infection and not enhanced replication. In further support of this, replication of WT rPOWV and rPOWV NS3 T561M was equivalent in *Ixodes scapularis* ISE6 cells suggesting no selective advantage in cells of a relevant tick vector (Figure 6D). Finally, pathogenesis of WT rPOWV and rPOWV T561M was compared in C57BL/6J mice. The closest TRIM5 ortholog in mice, Trim30d, inhibits replication of tick-borne orthoflaviviruses but by recognition of NS5 and not NS3 (*16*). WT POWV infection was uniformly lethal whereas 25% of mice survived infection with rPOWV NS3 T561M (Figure 6E). Weight loss curves were similar following infection with the two variants (Figure 6F) and no differences were observed in virus titers in the spleen or brain over the course of infection (Figure 6G, H), or in inflammatory pathology in the CNS (Supplemental Figure 1). Thus, the T561M mutation does not confer a replication advantage in models of enzootic hosts (ticks and rodents) that do not have a restrictive TRIM5α allele.

**Figure 6:**
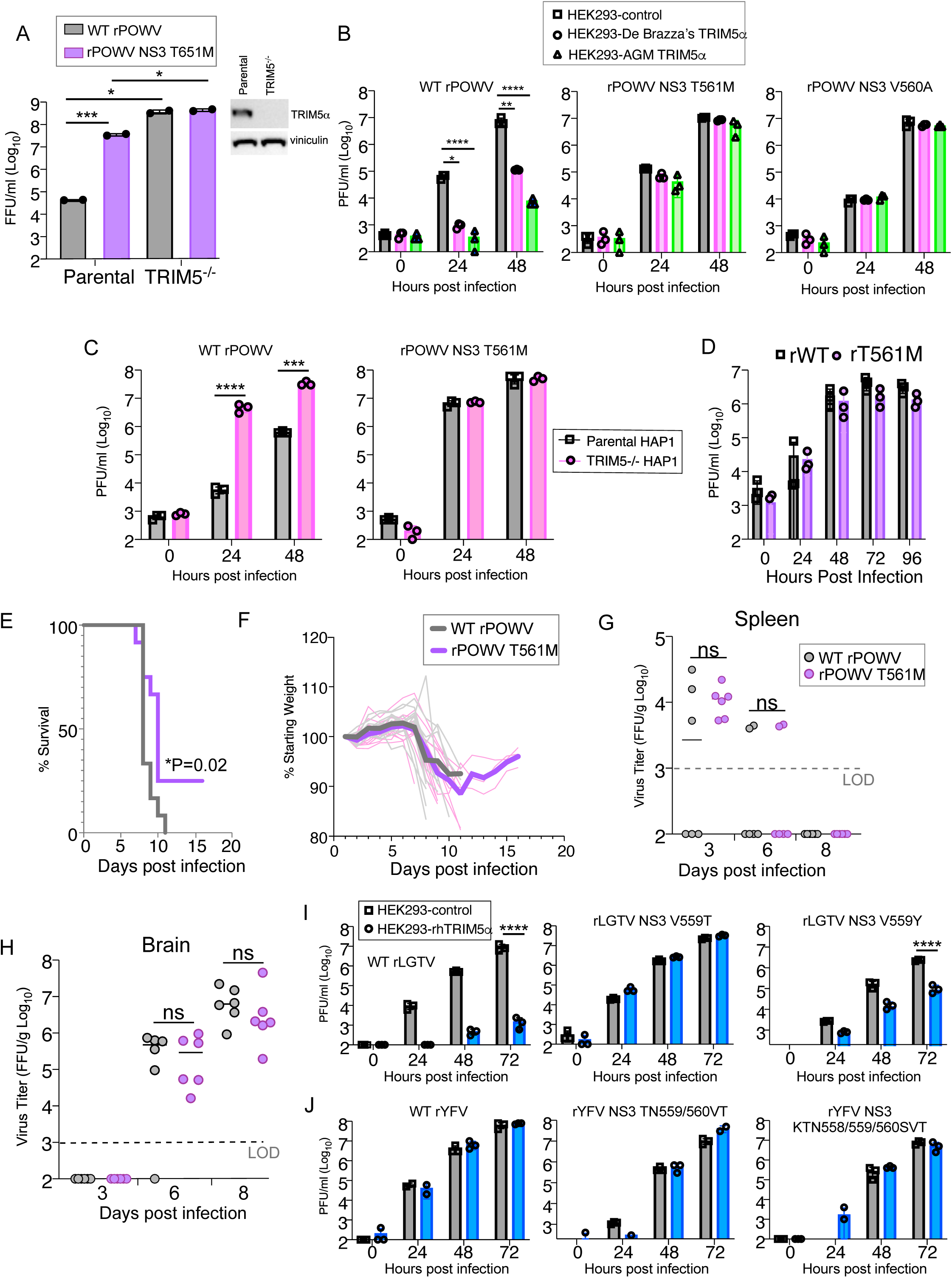
Powassan virus T561M has no virus-intrinsic replication advantage. **A.** Virus titers at 48 hpi of parental and TRIM5^-/-^ Vero cells (n=2). **B.** Growth curves of WT rPOWV or virus variants in HEK293 cells stably expressing De Brazza’s or AGM TRIM5α-HA or in control cells (n=3). **C.** Growth curves of WT rPOWV or virus variants in parental and TRIM5^-/-^ HAP1 cells (n=3). **D.** Growth curves of WT rPOW WT or NS3 T561M in ISE6 cells (MOI 0.1) and titered on BHK cells (n=3). **E.** Kaplan-Meier survival curves and **F.** Weight loss monitoring of C57Bl/6 mice infected with 1000 PFU WT rPOWV WT or NS3 T561M via the footpad (n=12/group). Virus titers in **G.** Spleen and **H.** Brain measured by focus forming units (FFU) on BHK cells (LOD=limit of detection; n=6). **I.** Virus titers in Growth curves of rLGTV bearing non-synonomous mutations at NS3 V559 present in mosquito-borne orthoflaviviruses in HEK293-rhTRIM5α-HA cells (n=3). **J.** Growth curves of rYFV bearing non-conserved substitutions implicated in TRIM5α resistance in HEK293-rhTRIM5α-HA cells (n=3). Graphed values represent the mean ± SD. Statistical significance was assessed using one-way ANOVA followed by Sidak’s post hoc test for multiple groups. ns: not significant, * p<0.05, ** p<0.01, *** p<0.001, **** p<0.0001.

Given the common ability of distinct tick-borne orthoflaviviruses to avoid TRIM5α-mediated restriction through substitution at V559, we determined the effect of V559 mutation to equivalent residues found in mosquito-borne orthoflaviviruses (Thr or Tyr; Figrue 5A). Interestingly, rLGTV encoding V559T (found in YFV) was highly replication competent and resistant to restriction by rhTRIM5α. rLGTV NS3 V559Y (present in DENV, WNV and JEV) produced 10-fold lower infectious virus but was more resistant to TRIM5α than WT rLGTV (Figure 6I). Given the tolerance of the YFV Thr residue here, we then tested if the tick-borne V559/T560 residues conferred TRIM5α sensititivity to YFV as a naturally resistant virus. However, substitutions in NS3 at TN559/560VT or KTN558/559/560SVT rendered rYFV partially replication defective and retained insensitivity to rhTRIM5α when compared to WT rYFV (Figure 6J). Thus, as expected, orthoflavivirus sensitivity to the antiviral function of TRIM5α cannot be attributed to a single substitution or minimal linear motif in the NS3 protein and is instead influenced by multiple determinants.

### Replication of POWV NS3 T561M in human MDDCs induces rapid and heightened IFN and cytokine/chemokine responses

Identification of a naturally occuring tissue-culture adaptation of POWV to evade TRIM5α provides the tools needed to examine how TRIM5α functions in primate immunity to tick-borne orthoflaviviruses. Infection of human monocyte-derived dendritic cells (MDDCs) from 5 donors with rPOWV revealed that TRIM5α reduces replication of POWV by at least 100 fold in the first 24 h of infection. However, replication of WT rPOWV increased over the next 24 h while rPOWV NS3 T561M titers decreased (Figure 7A). Cytokine and chemokine expression was generally not evident until 48 hpi with WT rPOWV, whereas moderate-to-high levels of cytokines were expressed by rPOWV NS3 T561M-infected cells at 24 hpi including IFNβ, IFNγ, CXCL9, CXCL10, IL-6, TNF, IL-8, CCL2 and CCL5, and further increased by 48 hpi (Figure 7B, C). Differences in POWV genomic RNA levels were observed at 24 hpi, concomitant with induction of *IFNB1* and *CXCL10* mRNA, but not at earlier time points of 6 and 12 hpi (Figure 7D, E, F). RNAseq performed on the same 5 donor MDDCs at 24 hpi revealed that the top differentially expressed genes (DEGs) were classical antiviral interferon stimulated genes (ISGs) with demonstrated anti-orthoflavivirus function (*17–21*), including RSAD2 (viperin), IFIT1, IFIT2, IFIT3, CMPK2, IFIH1 and OASL (Figure 8A). Further, the top 10 pathways differentially upregulated by rPOWV NS3:T561M were driven by interferon and interleukin signaling, while the top 10 downregulated pathways were related to cellular translation, amino acid metabolism, mitochondrial translation, TCA cycle and electron transport, consistent with increased virus burden (Figure 8B). Inhibition of JAK-STAT signaling using the JAK inhibitor Ruxolitnib reduced amplification of IFNβ expression while increasing rPOWV NS3:T561M replication, but did not change viral RNA amplification in MDDCs infected with WT rPOWV (Figure 8 C,D). Thus, high early IFN signaling is limiting to the replication of TRIM5α-resistant POWV, but not WT virus. These data demonstrate that TRIM5α is a significant barrier to replication of POWV in human DCs. However, while TRIM5α is clearly an important restriction factor in this context, POWV replication in the absence of TRIM5α-mediated restriction generates an early, robust antiviral and inflammatory response that is likely to have negative consequences for virus dissemination through establishment of the antiviral state in tissues and mobilization of adaptive immunity (*22–24*). Together these data suggest that TRIM5α is a potent intrinsic cellular barrier to infection with tick-borne orthoflaviviruses that restrains replication to levels that aid the virus in avoiding innate immune recognition (Figure 9).

**Figure 7:**
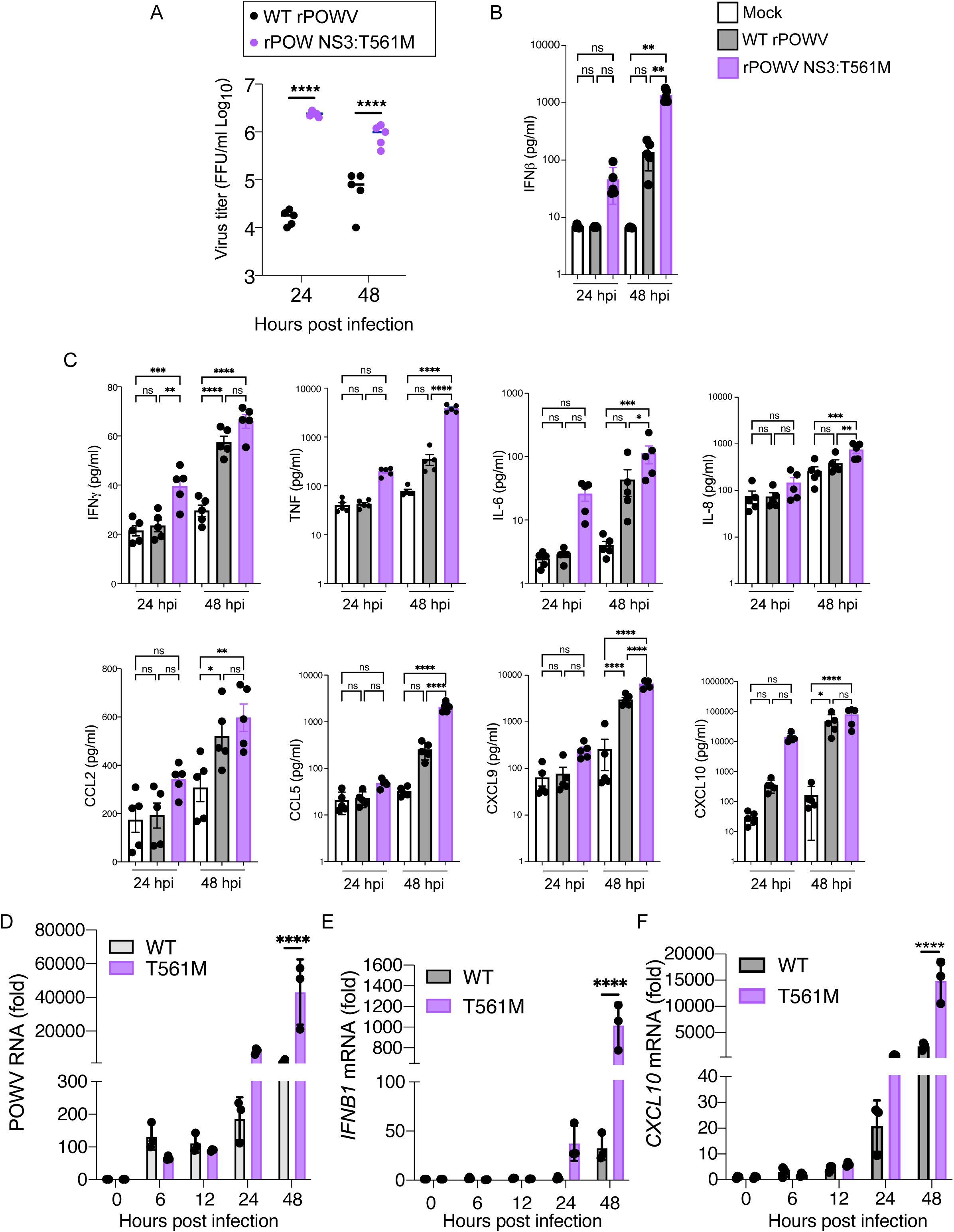
Replication of POWV NS3 T561M in human MDDCs induces rapid and heightened IFN and cytokine/chemokine responses. **A.** Growth curves of WT rPOWV and rPOWV NS3 T561M in MDDCs from 5 human donors. **B.** IFNβ protein expression measured by ELISA. **C.** Cytokine and chemokine concentrations in cell supernatants measured by Bioplex assay. **D.** Quantification of POWV positive sense RNA, **E.** *IFNB1* mRNA or **F.** *CXCL10* mRNA in MDDCs from 3 human donors. Graphed values represent the mean ± SD. Statistical significance was assessed using one-way ANOVA followed by Sidak’s post hoc test for multiple groups. * p<0.05, ** p<0.01, *** p<0.001, **** p<0.0001.

**Figure 8:**
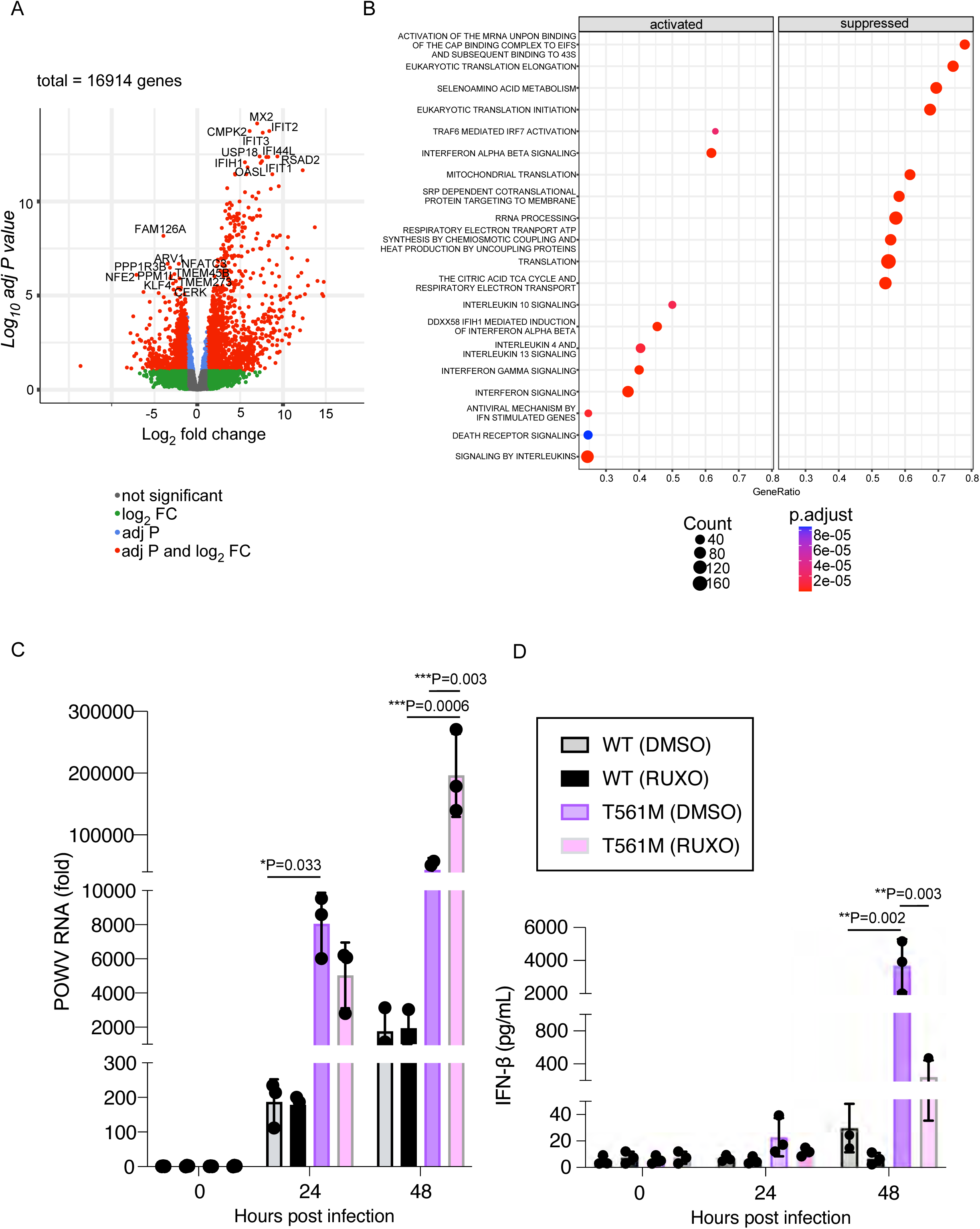
Replication of POWV NS3 T561M in human MDDCs induces heightened antiviral responses. **A.** Volcano plot of differentially expressed genes, and **B.** Reactome pathway analysis comparing differences following infection with WT rPOWV and rPOWV NS3 T561M at 24 hpi (n=5). **C.** Quantification of POWV positive sense RNA, and **D.** IFNβ protein expression measured by ELISA, in MDDCs pre-treated with Ruxolitnib and infected with POWV (n=3). Graphed values represent the mean ± SD. Statistical significance was assessed using one-way ANOVA followed by Sidak’s post hoc test for multiple groups. * p<0.05, ** p<0.01, *** p<0.001, **** p<0.0001.

**Figure 9:**
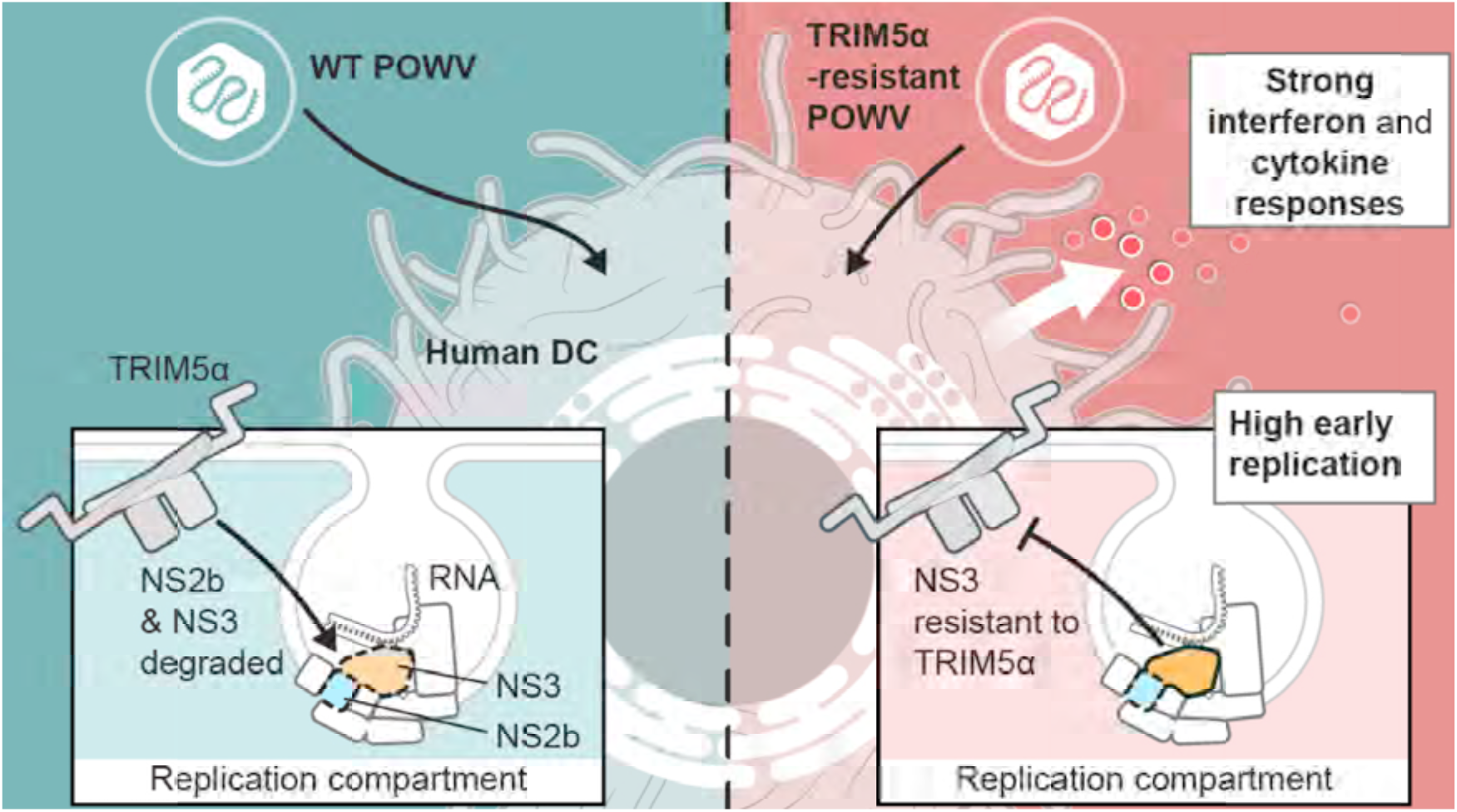
Schematic overview of TRIM5α-mediated restriction of POWV replication. TRIM5α is a significant barrier to replication of POWV in human DCs by binding to the viral NS3 protein. POWV escape from TRIM5α enables high virus replication and generates an early, robust antiviral and inflammatory response that is likely to have negative consequences for virus dissemination through establishment of the antiviral state in tissues and mobilization of adaptive immuninity. Thus, TRIM5α restrains replication of tick-borne orthoflaviviruses to levels that aid the virus in avoiding innate immune recognition.

## Discussion

Rhesus macaque TRIM5α was first reported as a consequential cell-intrinsic inhibitor of HIV-1 in 2004 (*5*), with subsequent analysis of structure and function, combined with evolutionary patterns of *TRIM5* in primates, indicating that a primary function of TRIM5α is to protect against retroviruses and retrovirus elements (*4, 8*). However, recent reports have demonstrated that primate TRIM5α has antiviral activity against orthoflaviviruses and orthopoxviruses, demonstrating a broad antiviral potential (*11, 12, 25*). TRIM5α must therefore encode flexibility in recognition of diverse viral molecular patterns (*26*). Further, TRIM5α appears to have multiple effector functions to engage different antiviral pathways in the cell, including immune activation and protein degradation pathways, dependent on the invading virus (*6, 12, 27–31*). Here we have begun to apply the knowledge and research approaches that have defined salient features of primate TRIM5α as a cell-intrinsic barrier to retrovirus replication in the context of orthoflaviviruses.

The TRIM5α PRYSPRY domain is critically required for restriction of orthoflaviviruses as the domain required for substrate binding to NS3 (*11, 12*). By testing a panel of old world and new world TRIM5α variants, we found that restriction appeared limited to old-world species. Interestingly, TRIM5α from *C. aethiops* and *C. sabaeus* shares 97.8% identity at the amino acid level, with 6 of the 8 non-synonomous differences occurring with the first variable domain within the PRYSPRY. Within the *TRIM5* PRYSPRY domain, the majority of positively selected sites are clustered within four variable domains (V1 to V4) that are in direct contact with retroviral capsids (*8, 32–38*). Based on analysis of the highly flexible V1 loop that contains the principal determinants for capsid-binding specifity, *TRIM5* in the Papionini (which includes macaques, baboons and mangabeys) and Cercopithecini tribes (which includes African Green monkeys, patas monkeys and guenons) is thought to have evolved independently to restrict endemic lentiviruses through recognition of common or closely overlapping sites on the capsid protein (*33*). Thus, TRIM5α specificity-determining sequences recognizing lentiviral capsids may also be involved in NS3 binding. This conclusion remains to be experimentally addressed but is supported by our observations that i) greater antiviral function of *C. aethiops* TRIM5α compared to the closely related *C. sabaeus*, and ii) that TRIM5α sequences from both Papionini and Cercopithecini species restricted replication of LGTV and TBEV. Our initial characterization of restriction potential of different primate TRIM5α sequences towards orthoflaviviruses provides the toolbox for swapping of variable loop sequences mirroring the classical studies performed in the retrovirus field to identify sequence determinants of TRIM5α-dependent restriction.

Orthoflavivirus evasion of TRIM5α was achieved through a single non-synonomous substitution in a flexible loop that connects domain II and III of the NS3 RNA helicase. NS3 functions in the replication organelle to unwind dsRNA during RNA replication and to provide single-stranded RNA for packaging into virions (*39, 40*). Here, NS3 is in complex with NS5 to feed newly unwound negative-sense template strand to the NS5 RNA-dependent RNA polymerase for synthesis of the positive-sense genomic RNA (*40*). The NS3 helicase first interacts with the double-stranded replicative form at the NS3 helical gate and β-wedge, a structural region that spans subdomains II and III (*40*). Therefore the V559 loop may regulate positioning of subdomain III in relation to subdomain II. We did not observe increases in rPOWV NS3:T561M RNA until after 12 hpi in MDDCs, suggesting that TRIM5α restriction occurs late in the infectious cycle, and may target roles of NS3 in coupling RNA replication to virion assembly. Future work using viral replicons will further define the precise events sensitive to restriction. Interestingly, the same solution to restriction was achieved by LGTV (at V559) and POWV (at T561) suggesting a conserved mechanism of recognition by TRIM5α.

The precise molecular interactions between PRYSPRY and retrovirus capsids have been challenging to capture and different binding models have been proposed (*4*). For example, studies suggest that the PRYSPRY domain of a single TRIM5α molecule engages in interhexamer binding of capsid protein, such that the V1 loop interacts with one capsid hexamer while the V2 and V3 domains interact with an adjacent capsid hexamer (*4, 41–44*). However, most Old World primate TRIM5α proteins restrict the lentivirus HIV-1 and the gammaretrovirus N-tropic MLV despite very little sequence homology (*45*). The lack of capsid-sequence specificity suggests that PRYSPRY domains exibit high plasticity in binding (*4, 26*). TRIM5α also uses plasticity in the second B-box domain and coiled-coil domains to form dimers (the fundamental oligomeric state) or trimers to achieve templated assembly to bind to the incoming mature virus core consisting of approximately 250 capsid hexamers (*46–48*). This templated assembly overcomes low binding affinity of PRYSPRY for individual capsid hexamers, and facilitates trimeric assembly of TRIM5α RING E3 ligase domains for activation of innate immune signaling (*49, 50*). It is difficult to speculate without further experimentation how TRIM5α PRYSPRY recognizes the orthoflavivirus NS3 protein and whether higher-order TRIM5α assembly is required for restriction. However, we think that a minimum of TRIM5α dimers are important as expression of a RING E3 ubiquitin ligase mutant of TRIM5α stabilized NS3 following co-expression in human HEK293 cells (*12*), indicating a dominant negative function of the ectopically expressed mutant.

Additional evidence suggests that TRIM5α targets NS3 in the replication organelle including that i) resistance of LGTV to TRIM5α may have involved selection of viral quasispecies sequences in LGTV NS5, ii) localization of TRIM5α to the replication organelle by TEM in LGTV-infected cells, and iii) loss of TRIM5α co-localization with the viral replicative intermediate dsRNA associated with viral evasion for both LGTV and POWV. However, our previous interaction mapping studies demonstrated that the NS3 helicase domain was not required for TRIM5α binding (*12*). Instead, the protease domain together with the cytoplasmic peptide of NS2B that enables assembly of the protease active site, and the short linker domain between the protease and helicase domains of NS3 (up to residue 190) were all required to observe degradation of NS3 by TRIM5α (*12*). Within the replication organelle, the role of NS2B is to anchor NS3 and NS5 to the membrane and orient the NS3 helicase domain (*51*). The role of the protease is to cleave the viral polyprotein, which may take place on ER membranes adjacent to the replication organelle, and to cleave cellular substrates to evade intrinsic antiviral processes including host IFN and autophagy responses (*52, 53*). Although suggestive that the replication organelle is targeted by TRIM5α, our data do not fully discern if specific functions of NS2B/3 are targeted for host antagonism. Further, the observation that TRIM5α can still sense the infection status of the cell in the context of rPOWV NS3:T561M infection even though localization to dsRNA is compromised indicates that TRIM5α responds to stimuli more complex than a binary interaction. How TRIM5α accesses NS3 and the replication organelle that is otherwise highly shielded from recognition by pattern recognition receptors (*54, 55*) is a critical question that could inform therapeutic approaches to better alert the innate immune response to the infection threat.

Vero cells derived from African green monkey (also known as sabeus monkey) are commonly used to propagate virus stocks due to their inability to produce type I IFNs (*56*), and we used this cell line to generate working stocks of POWV. However, POWV replicates comparatively better in baby hamster kidney (BHK) cells that are also deficient in antiviral interferon responses (*57*), hinting at the presence of restriction factors that inhibit POWV replication in Vero cells. The NS3 T561M mutation in our POWV stocks was most likely naturally acquired to evade TRIM5α following repeated propagation in Vero cells. Interestingly, adaptive mutations in a POWV replicon grown in human Huh7 cells naturally occurred in NS2A R195K, NS3 G122G (synonymous substitution), and NS3 V560M (*58*), of which V560M is the same substitution we identified as sufficient for LGTV to escape TRIM5α, and we experimentally demonstrated to confer substantial resistance for POWV. Thus, primate TRIM5α is a strong natural barrier to infection with tick-borne orthoflaviviruses and may function in multiple cell types, including liver and kidney epithelial cells (Huh7 and Vero respectively) in addition to monocyte-derived cells. Roles for TRIM5α beyond DCs, monocytes, and T cells have not been explored due to the primary focus on HIV-1 and related viruses in this field. However, understanding the relevance of TRIM5α in a broader array of cell-types and tissues is important given the newly identified relevance of TRIM5α to unrelated viruses, including orthoflaviviruses and orthopoxviruses (*12, 25*).

Neuroinvasion is a relatively rare outcome of infection with POWV. Therefore, we looked for mutations at NS3 V560 or T561 in available virus sequences from 17 patients with POWV neuroinvasive disease (*59, 60*) as well as 599 sequences of both POWV lineage I and lineage II (deer tick virus) from *Ixodes* tick vectors available in the Virus Pathogen Resource (ViPR). However, based on consensus sequences, these mutations have not been observed. This suggests that NS3 may not be subject to TRIM5α-mediated evolutionary pressure in enzootic cycles. Experimentally, this conclusion is supported by equivalent replication of WT and TRIM5α-resistant POWV in *Ixodes* tick cells and lack of a replication advantage observed in a mouse pathogenesis model. Alternatively, viral resistance to TRIM5α may be actively selected against due to high activation of innate immune responses (see below). Together, the current available data suggest that escape from TRIM5α is not likely to be a primary determinant of neuroinvasion following POWV infection of humans.

The antiviral response is amplified following recognition of HIV-1 by TRIM5α by two mechanisms. First, trimers of the RING domain in the templated lattices of TRIM5α generate K63-linked ubiquitin chains that activate the TAK1 complex to drive AP-1 and NFκB signaling and upregulate interferon and cytokine expression (*6*). Second, TRIM5α-mediated destabilization of the capsid lattice enables sensing of reverse transcription products by cGAS-STING (*61*). In our studies, infection of MDDCs with rPOWV NS3:T561M resulted in higher production of infectious virus, and early, amplified antiviral and inflammatory responses compared to the WT virus. Restriction of WT POWV replication by TRIM5α was most strongly observed at 24 hpi when innate responses are relatively low. Thus, TRIM5α does not appear to contribute to the antiviral response of MDDCs infected with WT virus. However, the rPOWV NS3:T561M mutant does not completely evade TRIM5α, evidenced by recruitment of TRIM5α to perinuclear sites of infected cells and a ∼10-fold higher level of rPOWV NS3:T561M replication in TRIM5^-/-^ Vero cells compared to the parental cell line. Therefore we cannot conclude with confidence that enhanced IFN and cytokine/chemokine responses in infected MDDCs are TRIM5α-independent when high levels of replicative intermediates are being produced. Regardless, as suggested by our results using the JAK inhibitor Ruxolitnib, the high level of IFN and inflammatory response from infected MDDCs likely has negative consequences for virus amplification through establishment of the antiviral state. In vivo, this may reduce virus spread in tissues and mobilize adaptive immunity. Indeed, the related YFV live-attenuated vaccine strain, YFV-17D, replicates to very high titers that strongly drive IFN signaling, which is key to virus attenuation and vaccine efficacy (*2, 62–64*). Further, the highest DEGs induced in infected human MDDCs included genes known to significantly curtail replication of tick-borne orthoflaviviruses including RSAD2, IFIT1, IFIT2, IFIT3, CMPK2, IFIH1 and OASL (*17–19*). Thus, mutations in tick-borne orthoflaviviruses that escape TRIM5α may be selected against in primates to restrain replication to levels that avoid early activation of the host antiviral response.

## Acknowledgements

Thank you to Alexander Khromykh (University of Queensland) for providing the flavilinker sequence used in the generation of molecular clones for POWV, LGTV and YFV. Thank you to members of the Research Technology Branch of NIAID, Justin Lack and Paul Gardina for analysis of RNAseq data, and Ryan Kissinger for generation of the graphical abstract. This work was supported by the Division of Intramural Research, National Institutes of Allergy and Infectious Diseases, National Institutes of Health.

## Methods

### Ethics statement

Animal study protocols were reviewed and approved by the Institutional Animal Care and Use Committee (IACUC) at Rocky Mountain Laboratories (RML), NIAID, NIH in accordance with the recommendations in the Guide for the Care and Use of Laboratory Animals of the NIH. All animal experiments were performed in an animal biosafety level 3 (ABSL3) research facility at RML. Standard operating procedures for work with infectious POWV and protocols for virus inactivation were approved by the Institutional Biosafety Committee (IBC) and performed under BSL3 conditions. All work with viruses including recombinant work was reviewed prior to initiation by the Rocky Mountain Laboratories IBC and NIH Dual Use Research of Concern Institutional Review Entity (DURC-IRE), and the manuscript was reviewed prior to submission.

### Viruses

The viruses used in this study were handled at BSL2, BSL3, and BSL4 at the Rocky Mountain Laboratories according to approved institutional biosafety protocols. KFDV (P9605), TBEV (Sofjin), and POWV (LB) seed stocks were obtained from the University of Texas Medical Branch (UTMB). KFDV, TBEV, LGTV (strain TP21), and ZIKV (2013 French Polynesia)(*12*) seed stocks were propagated on Vero cells, and virus working stocks were aliquoted and frozen in liquid nitrogen tanks. The POWV lab stock was passaged 5 times in Vero cells and once in BHK-21 cells.

### Cell lines

Vero E6 cells (ATCC CCL-81), HEK293 cells (ATCC CRL-1573) , BHK-21 cells (ATCC CCL-10) and Hap1 (12) are commonly used experimental cells for propagating virus stocks and examining cellular responses. Cells were grown in Dulbecco’s modified Eagle media (DMEM) containing 10% fetal bovine serum (FBS) and 1% antibiotics Penicillin-Streptomycin in an incubator at 37°C and 5% CO_2_. ISE6 cells were kindly provided by Tim Kurtti (University of Minnesota) and grown in L15C300 media at 32°C. Cell lines were routinely tested for mycoplasma contamination.

### Generation of stable TRIM5α-expressing cell lines

TRIM5’s of different primate species were cloned into pWPI DEST lentivirus vectors with a C-terminal HA tag: chimpanzee (AY740617.1), De Brazza’s monkey (KP743978.1), grivet monkey derived from Cos-7 cells (AGMc) (AY843504.1), African green monkey (also known as sabeus monkey) derived from Vero cells (AGMv) (AY625003.1), patas monkey (AY843514.1), pygmy marmoset (AY843512.1), squirrel monkey (AY843517.1), red howler (AY843511.1), northern owl monkey (AY646198.1). Pigtailed macaque TRIMCyp sequence was amplified from *Macaca nemestrina* cDNA derived from PBMCs. The pigtailed macaque TRIMCyp sequence was most similar to DQ308405 with the following amino acid changes: K44E, E209K, T269A, H373R, S438G. Lentiviruses were generated by transfecting 5×10^6^ 293T cells with 1 µg pMD.G, 8 µg pSPAX2, and 8 µg pWPI vector using the ProFection Mammalian Transfection System (Promega). Supernatant was passed through a 0.45 micron filter and added to HEK293 cells. TRIM5 positive cells were selected with blasticidin S HCl at 5 µg/ml. Following construction of the stable TRIM5 cells, expression of each cell line was validated by western blotting with an HA antibody.

### Virus infections and plaque assay

For growth curve experiments, viruses were added to the plates at the indicated MOI. Cells were incubated with inoculum 1 hr followed by 1-2 washes with PBS. Fresh media was added to the cells, and supernatants were pulled at the indicated times and frozen at -80°C. Limiting dilution plaque assays were performed as previously described (12). Briefly, 10-fold dilutions of viral supernatant were plated on 48-well plates of Vero cells (LGTV, KFDV, TBEV, YFV) or BHK cells (POWV) followed by a 1.5% carboxymethyl cellulose overlay. When the plaques formed, plates were flooded with 10 % formalin. Cells were stained with 1 % crystal violet, and plaques were counted.

### Generation of LGTV, POWV, and YFV molecular clones

Molecular clones were generated using circular polymerase extension reaction (CPER) methods previously described for Zika virus (*14*). All work was reviewed by the NIAID Institutional Biosafety Committee and the Dual Use Research Committee (DURC). The molecular clones for LGTV-TP21, POWV-LB, and YFV-H196 were based on published sequences EU790644.1, L06436, and MF538784.1 respectively. For each virus, the genome was broken into 7 overlapping fragments that were between 500-2200 kb in length. These fragments along with a flavilinker fragment were synthesized in either pUC19 or pJet1.2. A polymerase chain reaction (PCR) was performed using Q5 high fidelity DNA polymerase (New England Biolabs) to amplify each fragment, and the PCR products were gel-purified using the QIAquick Gel Extraction Kit (Qiagen). See Supplemental Tables 1-3 for a list of primers used for the infectious clones. Finally, 0.1 pmol of each fragment was combined and a CPER PCR was performed using Q5 high fidelity DNA polymerase using CPER PCR conditions as described previously (*14*). The resulting PCR product was transfected directly into Vero cells or BHK cells using TransIT-LT1 transfection reagent (Mirus) or Lipofectamine LTX and PLUS Reagents (Thermofisher). CPE typically occurred within day 7-10 post transfection of the LGTV clone in Vero cells and within 3 days after transfection with the POWV and YFV clone in BHK cells. Recovered virus was sequence confirmed.

### Immunofluorescence

293 cells were plated on 4-well chamber slides and infected as indicated. Cells were fixed with 4% PFA for 30 minutes (LGTV) or overnight (POWV). Permeabilization was performed with 100% ice-cold methanol for 5-8 minutes. Cells were blocked with 3% normal goat serum (NGS), and labeled with primary antibodies overnight at 4°C in 1% NGS/PBS solution: anti-Flag (Cell Signalling, 86861) 1:200 and anti-dsRNA (Scicons) 1:400. Cells were washed 3-4 times with PBS, and secondary antibodies were added at 1:2000 dilution in 1% NGS/PBS for 1 h at room temperature. Cells were washed and mounted on coverslips with ProLong Gold Anti-fade reagent containing DAPI (ThermoFisher). Confocal images were obtained using a Zeiss LSM710 confocal microscope and analyzed using Zen software (Carl Zeiss).

### Apex staining and Transmission Electron Microscopy

HEK293 cells stably overexpressing rhesus TRIM5-Apex2 or mCherry-empty vector control cells were plated on Thermanox Coverslips (Electron Microscopy Sciences) and infected with LGTV-WT or LGTV-VA recombinant viruses for 24 h. Cells were washed 1x with PBS and fixed with 2% glutaraldehyde. For Apex labeling, the samples were incubated with SIGMAFAST DAB with Metal Enhancer tablets (Milipore Sigma) diluted in water for 8 min on ice. The DAB solution was removed and cells were incubated with 20mM glycine in PBS for 5 min.

### Antibodies

HA-tagged constructs for western blotting were detected using a 1:5,000 dilution of anti-HA-peroxidase antibody (Roche clone 3F10, #12013819001). HA-tagged constructs for indirect immunofluorescence were detected using anti-HA (Zymed, #71-5500). β-actin was also detected as a loading control using a 1:10,000 dilution of mouse anti-β-actin (Sigma, A5441). A 1:3,000 dilution of goat anti-mouse (Dako, #P0447), anti-rabbit (Thermo Scientific, #P0448) or anti-chicken (Millipore, #12-341) horseradish peroxidase-conjugated antibody was used as a secondary probe. Blots were developed using the ECL Plus detection reagent (GE Healthcare, #RPN2132). Antibodies to detect viral antigens, LGTV (NS3) (previously described in Taylor et al., 2011), and dsRNA antibody J2 (English& Scientific Consulting, #10010200), anti-DYKDDDDK rabbit mAb (D6W5B; Cell Signaling Technologies).

### Immunoprecipitation (IP) and Western Blot Analysis

HEK293 cells were washed three times with DPBS and lysed on ice in RIPA buffer (50 mM Tris-HCl [pH 7.6], 150 mM NaCl, 0.1% SDS, 1% Igepal, and 0.5% Na-deoxycholate) with protease inhibitor cocktail (Roche). For IPs of overexpressed proteins, 2 wells of a 6 well dish at 1×10^6^ cells/well were used per reaction; for IPs of virus-infected stable TRIM5 HEK293 cells, a 10cm dish of 7×10^6^ cells/dish was used per reaction; for detection of endogenous TRIM5, HEK293 or HAP1 cells were grown to confluency in 3-4 T150 tissue culture flasks. Samples were subjected to centrifugation for 10 min at maximum speed to remove cellular debris. Protein G-conjugated agarose beads (Roche) or PrecipHen for chicken antibodies (Aves Labs) were used to clear cell lysates at 4°C for 3 h. Samples were centrifuged to remove beads, and 2 μg of antibody analogous to the protein of interest was added to each lysate for 1 h with rotation at 4°C. 50 μL protein G-agarose or PrecipHen beads and were incubated with rotation at 4°C overnight. Lysates were subjected to centrifugation, and beads were washed three times with RIPA buffer prior to elution by incubation at 95°C in 1 × sample buffer (62.5 mM TRIS [pH 6.8], 10% glycerol, 15 mM EDTA, 4% 2-ME, 2% SDS, and bromophenol blue). For western blot analysis HEK293 cell lines were grown to confluency in a 12-well or 6-well dish, collected using a cell scraper, and lysed in RIPA buffer containing complete protease inhibitor (Roche, #11836170001). After quantification of protein concentration using a Bradford assay, 30 μg of whole cell extract was resolved using a 10% polyacrylamide gel and transferred to a nitrocellulose membrane.

### Mouse model

C57BL/6J (6 weeks old) mice (The Jackson Laboratories, 12 mice per group 6M, 6F) were inoculated under isoflurane anesthesia in the left hind footpad with 20 μL PBS containing 1000 PFU of either WT or T561M rPOWV. Mice were weighed daily and monitored for clinical signs of infection. Mice were euthanized when they met pre-determined endpoint criteria of 20% weight loss or signs of neurological infection, including tremor, seizure or limb paralysis.

### MDDCs

Human monocyte-derived dendritic cells were prepared as previously described [70]. Briefly, we obtained human monocytes from 5 different donors from the NIH Clinical Center, Department of Transfusion Medicine (deidentified), . Monocytes were frozen in RPMI with 40% FBS, 10% DMSO and stored in liquid nitrogen until needed. Thawed monocytes were washed with PBS and resuspended at 1×10^6^ cells/mL in DC medium [RPMI + Glutamax (Invitrogen), 5% FBS, 15 mM Hepes, 0.1 mM nonessential amino acids, 1 mM sodium pyruvate, 100 units/mL penicillin and 100 μg/mL streptomycin] containing IL-4 (20 ng/mL) and GM-CSF (20 ng/ml) (PeproTech) and cultured for 5d with replacement of half of the medium and addition of fresh cytokines every other day. Nonadherent DC were harvested by centrifugation and suspended in DC medium. The resulting cells were determined to be >95% CD11c+/CD209+ by flow cytometry. MDDCs were pretreated with Ruxolitnib (Invivogen; 10ug/ml) for 1 h prior to infection, and then kept in cultures for the duration of experiments.

### Bulk RNA-seq of human MDDCs

Human MDDCs were plated at 1×10^6^ cells/well in a 12 well format and mock-infected or infected with POWV at an MOI of 0.1 in a small volume for 1 h at 37°C, 5% CO_2_ with occasional gentle rocking. MDDCs were harvested by centrifugation, lysed with 1 mL of TRIzol reagent/sample (Thermo Fisher Scientific), and samples were stored at -80°C before extraction and bulk RNA-seq. Cell supernatants were collected for cytokine and viral plaque assays.

For RNA extraction, samples were mixed with 200 µL of 1-bromo-3-chloropropane (Sigma Aldrich) and centrifuged at 4°C for 15 minutes at 16,000 x g. The aqueous phase was removed and passed through a QIAshredder column (Qiagen) at 21,000 x g for 2 min. RNA was extracted using the Qiagen AllPrep DNA/RNA system. An additional on-column DNase I treatment was performed for all extractions. RNA quality was assessed using the Agilent 2100 Bioanalyzer RNA 6000 Pico kit (Agilent Technologies) and quantified using the Quant-it RiboGreen RNA assay (Thermo Fisher Scientific).

Following RNA extraction, 200 ng RNA was used as the input for the Illumina stranded mRNA prep, ligation kit (Illumina) to generate sequencing libraries following the manufacturer’s protocol. The final libraries were analyzed using the Agilent bioanalyzer and the libraries were quantified using the Kapa SYBR FAST Universal qPCR kit for Illumina sequencing (Kapa Biosystems). The individual libraries were diluted to 6 nM and 2 uL of each added to a library pool. The library pool was denatured and diluted to a 10 pM stock and paired-end 2 x 75 cycle sequencing carried out on the MiSeq using a Nano V2 flow cell and 300 cycle chemistry. Following the sequencing run, reads per microliter mapping to human genes were determined for each library. The library pool was rebalanced and quantified using the Kapa SYBR FAST Universal qPCR kit for Illumina sequencing. The library pool was diluted to 9 pM and paired-end 2 x 75 cycle sequencing carried out on the MiSeq using a Nano V2 flow cell and 300 cycle chemistry. The final library pool containing 6 nM of each library was sent to the National Cancer Institute, Center for Cancer Research Sequencing Facility (NCI CCR-SF) for further sequencing. The samples were paired-end 2 x 100 cycle sequenced using a NovaSeq 6000 instrument and SP flow cell and 300 cycle chemistry.

Following sequencing, Raw fastq files were trimmed to remove adapters and low-quality bases using Cutadapt v1.18 before alignment to the GRCh38 reference genome and the Gencode v42 genome annotation using STAR v2.7.9a (*65*). PCR duplicates were marked using the MarkDuplicates tool from the Picard v3.1.0 (https://broadinstitute.github.io/picard/) software suite. Raw gene counts were generated using RSEM v1.3.3 (*66*) and were filtered to include genes with >=1 count per million (CPM) in at least 2 samples.

Differential expression was evaluated using limma with TMM normalization (*67*). For all differential expression comparisons, pre-ranked geneset enrichment analysis (GSEA) (*68*) was performed using the Reactome Database (*69*).

## Statistics and Reproducibility

All graphical values are shown as mean ± standard deviation. Statistical analysis was performed using GraphPad Prism 10. Ordinary one-way or two-way ANOVAs with Dunnett’s or Tukey multiple comparisons tests were performed, as indicated in figure legends, to test statistical significance between means. P-values less than 0.05 were considered statistically significant. Data are derived from a minimum of three replicates. Animal studies were conducted with an N of 12 mice per group for surviviral and 6 mice per group for quantification of virus in tissues.

## Data Availability Statement

All data supporting the findings of this study are available within the paper and its Supplementary Information, with the exception of RNA-Seq data which was deposited into the Gene Expression Omnibus database under accession number GSEXXX and are available at the following URL: XXXX.

