## Supplemental Tables 1-3 for "Restraint of Powassan virus replication by TRIM5α facilitates viral avoidance of antiviral immunity"

**Table 1:** Primers used to generate LGTV-TP21 molecular clones by CPER

| **LGTV TP21 CPER primers** | **Forward Primer (5’-3’)** | **Reverse Primer (5’-3’)** |
| --- | --- | --- |
| Fragment 1 | AGATTTTCTTGCGCGTGCATGCGT | CGCTGTCTGTGGGTGCTCTTTTCC |
| Fragment 2 | GGAAAAGAGCACCCACAGACAGCG | CCTTCACTGGAATGCCCATCTCCAAC |
| Fragment 3 | GTTGGAGATGGGCATTCCAGTGAAGG | CAAATGACACTCTTCCCTCGCTGCC |
| Fragment 4 | GGCAGCGAGGGAAGAGTGTCATTTG | ACGGCCCTCTCTAAACATACGGGC |
| Fragment 5 | GCCCGTATGTTTAGAGAGGGCCGT | CAGGGTGTCTCCCTCGGATCC |
| Fragment 6 | GGATCCGAGGGAGACACCCTG | CCATCATCCTCACCAGCTGCACC |
| Fragment 7 | GGTGCAGCTGGTGAGGATGATGG | AGCGGGTGTTTTTCCGAGACACG |
| Flavilinker | GTGTCTCGGAAAAACACCCGCTGGGTCGGCATGGCATCTCCACC | GCATGCACGCGCAAGAAAATCTCGGTTCACTAAACGAGCTCTGCTTATATAG |

**Table 2:** Primers used to generate POWV-LB molecular clones by CPER

| **POWV LB CPER primers** | **Forward Primer (5’-3’)** | **Reverse Primer (5’-3’)** |
| --- | --- | --- |
| Fragment 1 | AGATTTTCTTGCACGTGTGTGCGGGTGCT | GATCCGACATGGCTTGTCGCTTCCTGT |
| Fragment 2 | ACAGGAAGCGACAAGCCATGTCGGATC | CTTGATTATGAGCAACCCCATGGCCAGTGC |
| Fragment 3 | GCACTGGCCATGGGGTTGCTCATAATCAAG | CTATGGCGCCTCCCTTGGCTATCGATG |
| Fragment 4 | CATCGATAGCCAAGGGAGGCGCCATAG | GTCCAGTCAGGATGTCCACGGCACT |
| Fragment 5 | AGTGCCGTGGACATCCTGACTGGAC | CAAGCTAGTTTGGCACAGCCTCGCGATAC |
| Fragment 6 | GTATCGCGAGGCTGTGCCAAACTAGCTTG | CAGGAAGTACAGCGCCTTTCCGAACCTG |
| Fragment 7 | CAGGTTCGGAAAGGCGCTGTACTTCCTG | AGCGGGTGTTTTTCCGAGTCACACACCA |
| Flavilinker | GTGACTCGGAAAAACACCCGCTGGGTCGGCATGGCATCTCCACC | GCACACACGTGCAAGAAAATCTCGGTTCACTAAACGAGCTCTGCTTATATAG |

**Table 3:** Primers used to generate YFV-H196 molecular clones by CPER

| **YFV CPER primers** | **Forward Primer (5’-3’)** | **Reverse Primer (5’-3’)** |
| --- | --- | --- |
| Fragment 1 | AGTAAATCCTGTGTGCTAATTGAGGTGC | CCTTTTGGCACCTTCACTTGCATC |
| Fragment 2 | GATGCAAGTGAAGGTGCCAAAAGG | ACTGCGTTCAGGTACTTCCACAGA |
| Fragment 3 | TCTGTGGAAGTACCTGAACGCAGT | GCAGACATGATGCGCATTGTCTTC |
| Fragment 4 | GAAGACAATGCGCATCATGTCTGC | GAGTGCAAGAGCACGCTGATAGTA |
| Fragment 5 | TACTATCAGCGTGCTCTTGCACTC | CAATTTTGCGGTCCCTCTCGAGAC |
| Fragment 6 | GTCTCGAGAGGGACCGCAAAATTG | TCCGCCATCCTAATCAGTTGAACC |
| Fragment 7 | GGTTCAACTGATTAGGATGGCGGA | CCTTTGGATGACAAACACAAAACCACT |
| Flavilinker | CCTTTGGATGACAAACACAAAACCACTGGGTCGGCATGGCATCTCCACC | GCACCTCAATTAGCACACAGGATTTACTCGGTTCACTAAACGAGCTCTGCTTATATAG |
